# Generation of high-resolution *a priori* Y-chromosome phylogenies using “next-generation” sequencing data

**DOI:** 10.1101/000802

**Authors:** Gregory R. Magoon, Raymond H. Banks, Christian Rottensteiner, Bonnie E. Schrack, Vincent O. Tilroe, Terry Robb, Andrew J. Grierson

## Abstract

An approach for generating high-resolution *a priori* maximum parsimony Y-chromosome (“chrY”) phylogenies based on SNP and small INDEL variant data from massively-parallel short-read (“next-generation”) sequencing data is described; the tree-generation methodology produces annotations localizing mutations to individual branches of the tree, along with indications of mutation placement uncertainty in cases for which “no-calls” (through lack of mapped reads or otherwise) at particular sites precludes precise phylogenetic placement of mutations. The approach leverages careful variant site filtering and a novel iterative reweighting procedure to generate high-accuracy trees while considering variants in regions of chrY that had previously been excluded from analyses based on short-read sequencing data. It is argued that the proposed approach is also superior to previous region-based filtering approaches in that it adapts to the quality of the underlying data and will automatically allow the scope of sites considered to expand as the underlying data quality improves (e.g. through longer read lengths). Key related issues, including calling of genotypes for the hemizygous chrY, reliability of variant results, read mismappings and “heterozygous” genotype calls, and the mutational stability of different variants are discussed and taken into account. The methodology is demonstrated through application to a dataset consisting of 1292 male samples from diverse populations and haplogroups, with the majority coming from low-coverage sequencing by the 1000 Genomes Project. Application of the tree-generation approach to these data produces a tree involving over 120,000 chrY variant sites (about 45,000 sites if “singletons” are excluded). The utility of this approach in refining the Y-chromosome phylogenetic tree is demonstrated by examining results for several haplogroups. The results indicate a number of new branches on the Y-chromosome phylogenetic tree, many of them subdividing known branches, but also including some that inform the presence of additional levels along the “trunk” of the tree. Finally, opportunities for extensions of this phylogenetic analysis approach to other types of genetic data are noted.

## Introduction

Dramatic cost reductions and technological developments in massively-parallel short-read (or “next-generation”) sequencing have increased its use in human genetic studies in recent years. The data obtained from these studies are presenting the opportunity for a wide range of investigations involving the human Y chromosome. In contrast to alternative approaches such as microarray-based genotyping and Sanger sequencing, short-read sequencing provides an efficient means to probe large regions of the Y chromosome. The data from these sequencing studies carry enormous potential in the fields of genetic anthropology and genetic genealogy, allowing identification of novel phylogenetic markers and allowing significant refinement of the human Y chromosome phylogeny; for example, it has been shown that early data from the 1000 Genomes Project allow significant refinement of the R1b-M269 portion of the Y-tree based on 135 male samples [1]. The data also present the opportunity for improving the precision of haplogroup age estimates [2-4].

However, there are several challenges that must be overcome before data from such studies may be efficiently utilized to their full potential. Previous studies have explicitly [5] and implicitly [6] called attention to the challenges involved in using short-read sequencing data for large-scale Y-chromosome phylogeny construction. Issues with data quality (including the tendency for the data to suggest existence of false-positive variants), the potential for lack of mapped coverage at particular sites in individual samples (particularly in low-coverage datasets), and the sheer volume of data appear to be chief among these challenges.

Here, a phylogenetic analysis approach designed to work around these challenges is described.

## Description of the approach

### Variant calling

We employ diploid multi-sample variant calling through use of “mpileup” in the *samtools* software package [7]. In contrast to other approaches which rely on a set of single-sample variant calling results [8], this approach allows one to distinguish between results indicating a reference allele, and results indicating a lack of mapped coverage at individual sites in individual samples. (Results of single-sample variant calling typically only report sites that differ from reference allele, which can be misleading and can confound downstream results analysis, particularly for low-coverage sequence data.) These issues also reinforce the importance of having access to the raw mapped alignment data (e.g. BAM file) rather than a derived analysis like variant call data. Although not addressed or discussed in detail here, a noteworthy challenge with constructing a comprehensive multi-sample variant calling database can arise when datasets of multiple types must be merged (e.g. with mapped data aligned to different reference sequences or variant call data from Complete Genomics analysis pipeline), which can become a non-trivial bioinformatics task.

The details of the samtools variant calling command used here are presented in the Supporting Information.

Although it is tempting to employ haploid variant calling for the hemizygous Y-chromosome, we have found diploid calling to be more useful; the use of diploid calling admits the possibility for a (spurious) heterozygous result, and will return a genotype likelihood for such a result in each sample for each variant site. In addition to readily identifying suspicious results and potentially spurious variants, we find this information helpful for identifying sites that are prone to read mismapping or sequencing errors, as will be detailed later. Haploid chrY variant calling approaches, such as used in the Complete Genomics pipeline [9, 10], which omit this key information, make tree generation with an approach like that described here more challenging and error-prone.

The variant call file (VCF) produced through this analysis becomes the input to the next (genotype refinement) step.

### Genotype refinement

We next refine the genotype (GT) calls in the multi-sample variant call file based on genotype likelihoods (PL values). These refinements replace the original “0/0”, “0/1”, and “1/1” genotypes called by samtools with calls appropriate for the hemizygous Y-chromosome. In particular, “.” is incorporated to indicate “no-call”, “./.” is used to indicate a (spurious) heterozygous result, 0 is used to indicate reference result, and 1, 2, etc. are used to indicate results with an alternate allele. With the exception of the “./.” call, these calls conform to VCF specification. The details of how these “refined” genotypes are determined based on genotype likelihoods are presented in the Supporting Information.

### Variant filtering

There is a strong propensity for variant discovery analyses of short-read sequencing results to suggest the existence of false-positive variants, presumably due to mismapping of reads during the alignment process; this problem is particularly acute for the Y chromosome which has significant repetitive regions and regions with homology to other parts of the genome, such as the X chromosome [11, 12].

This propensity for mismapping will vary with details such as local sequence properties and read length, and certain regions of the Y chromosome are much more prone to this issue of false positives than others. Mismappings can similarly give rise to false negatives with low-coverage results. The challenge, then, in generating a Y chromosome phylogeny, is to avoid issues arising from spurious results at these unreliable sites. A common approach is to proactively remove these sites from consideration through a filtering step before subsequent phylogenetic analysis.

Previous studies of the Y-chromosome have employed position-based filtering, excluding regions known to be repetitive or have a high degree of sequence homology to other parts of the genome. Our investigations suggest that reliable results may be frequently obtained in the excluded regions of one widely-used position-based filter (encompassing about 9 Mb) [4] with phylogenetically informative variants being omitted from consideration in many cases. (For example, U106, a key SNP marker subdividing the R1b haplogroup, which is typically reliably genotyped with short-read sequence data, would be excluded by this prominent region-based filter.) Furthermore, even if the reliable regions are carefully and precisely determined for a given dataset, there is limited opportunity to generalize to other datasets, since the regions with reliable results will vary with methodological details (e.g. read length, the use of paired-end reads, and mapping algorithm, etc.). Here, we instead advocate for a filtering approach based on various properties of the underlying data, with the goal of implicitly adapting to methodological differences. The filtering described in this section serves as a “first-pass” with iterative reweighting serving to provide further refinement.

We note that spurious heterozygosity can be a key indicator of sites with unreliable results on the hemizygous Y chromosome. Heterozygosity will be less apparent from low-coverage results, but, considering enough low-coverage samples, these issues should become more evident. When considering these spurious heterozygous results, two particular modes of mismapping are considered here. The first, mismapping of reads with a non-reference allele at a site of interest, gives rise to false-positive variants. The other mismapping error mode is for reads with the reference allele to be mismapped to a site of interest, which can give rise to false-negatives. Thus, considering a number of samples simultaneously, filtering to remove sites with a high proportion of samples with heterozygous results might be considered to guard against the first mode of read mismapping. Similarly, filtering to remove sites with a high ratio of the number of samples with heterozygous results to the number of samples with “pure” non-reference allele result might be considered to guard against the second issue.

Additional indicators, commonly used in more general analyses of short-read sequencing data, such as strand-bias and mapping quality, may also be used as the basis for filtering results.

Thus, we use here many of the built-in filters implemented in the widely-used *vcf-annotate* tool in the *vcftools* package [13]. We have also implemented custom filters to use information such as heterozygosity. The details of the variant filtering approach used here are presented in the Supporting Information.

Additional possible filtering approaches, such as using the Y (mis)mapping pattern and/or variant results for a set of female samples, are possible, but have not been explored here.

Here, we also employ a temporary filtering to remove “singletons” (i.e. non-reference variants that appear to be present in only one sample within the dataset) to make the main tree generation steps more computationally efficient. We note that these variants can be readily localized to the terminal branches of the tree (leading to individual samples) *post facto* (e.g. during post-processing), without the need to consider them during the main tree generation process.

### Maximum parsimony tree generation

We use a maximum parsimony tree generation algorithm, as implemented in PHYLIP 3.69 [14], for individual tree generation steps. A key aspect of the chosen tree generation methodology is the ability to perform ancestral state reconstruction (i.e. localization of individual mutations to particular branches in the tree, to the extent supported by the underlying data). A modification to PHYLIP 3.69 source code was made by the authors to improve memory management for the ancestral state reconstruction, allowing for this critical analysis to be performed with reasonable memory requirements for the large datasets of interest here. The details of the maximum parsimony tree generation approach used here are presented in the Supporting Information.

### Iterative reweighting

The maximum parsimony tree generation algorithm readily allows for a weighting of individual variant sites in parsimony evaluation. We have developed a procedure of iterative reweighting of variant sites designed to downweight sites that are unstable (i.e. those that have a relatively high *in vivo* mutation rate and thus cannot reliably serve as “unique-event polymorphisms”) and/or are prone to unreliable results (due to, for example, read mismapping).

The idea is to use the number of mutations at a particular site in the results of a previous iteration of tree generation to inform the weight of the site in the next tree generation iteration. In particular, sites with higher mutation counts (generally indicative of instability or unreliability issues) are given reduced weight.

Here, we employ four iterations of reweighting following an initial, unweighted tree generation step. The first three iterations employ progressively stronger reweighting, while the final iteration also applies a proximity filter which reduces the weight of nearby variants appearing on the same branch to zero. The use of more aggressive reweighting in later steps is designed to allow the tree generation to proceed flexibly on a relatively flat “parsimony surface” in the early stages before the strong reweighting of later steps effectively “locks” the tree into place (similar to the annealing concept in the simulated annealing approach to global optimization [15, 16]).

The details of the iterative reweighting approach used here are presented in the Supporting Information.

### Post-processing of tree-construction results

The final step of the tree generation process is post-processing of the results from the last iteration of tree generation, to generate a text-based report suitable for human analysis and interpretation that details the mutations on each of the branches of the generated tree. The ancestral state table produced by PHYLIP is parsed to localize mutations, along with their polarity (i.e. whether a REF>ALT mutation or an ALT>REF mutation) to individual branches of the tree. The associated variant information (position, reference allele, and non-reference/alternate allele) are pulled in from the variant calling database. Additionally, the total number of mutations in the tree for each site are counted and reported to aid in manual interpretation of the results. Cases where the location of a variant is uncertain due to no-calls in key samples are also annotated to assist in the interpretation of results. (It is noted that uncertainty due to the existence of other trees with equivalent or similar parsimony are not considered here and such considerations would need to be made manually with the present approach.) This post-processing step also involves incorporation of the “singleton” variants into terminal branches; as discussed previously, these variants had been omitted from the tree-generation process for computational efficiency. Each of the mutations is also annotated if a match (either based on position, ancestral allele, and derived allele or based only on position) is found in a reference database of named chrY markers. [17]

In the process of analyzing the final results from this tree construction approach, we suggest that special scrutiny should be reserved for branches with a low number (*e.g.* fewer than three) of “solid” mutations (i.e. those mutations that do not carry uncertainty in position due to no-call and which have a low number of mutations throughout the tree); although these cases could very well be genuine, there is potential for errors in the underlying data or false positives to introduce spurious structure into the generated tree. We have also observed that different sequencing platform pipelines (e.g. SOLiD) can lead to platform-dependent artifacts (presumably false positive variants), which could adversely impact the generated tree structure in certain situations. Also, of course, as was implicit in the prior discussion on reweighting, sites with mutations at more than two or three branches in the phylogeny are likely to be too unstable and/or unreliable to serve as robust phylogenetic markers.

## Demonstration of the approach through application to a 1292-sample dataset

The approach described above is demonstrated through application to a dataset consisting of 1292 male samples from diverse populations and haplogroups. The majority of the sample data come from low-coverage sequencing by the 1000 Genomes Project [18]; the set of data from the 1000 Genomes Project also includes one high coverage sample (NA12891). Additionally, high-coverage data for ten male Human Genome Diversity Project (HGDP) samples and a Dinka sample, produced during a study by Meyer *et al.* [19], were incorporated into the analysis. Finally, low-coverage data for NA21313 (ERS037274), sequenced in a study by Wei *et al*. [4], were also incorporated. The source BAM filenames for the 1000 Genomes Project samples, indicating data/alignment details, are included in Supporting Information C. The University of Sheffield high performance computing cluster was used for all computationally-intensive analysis, including variant calling and maximum-parsimony tree generation.

An overview of the generated tree structure is presented in Figure 1. The raw results of the approach described here are presented in Supporting Information B and a figure depicting the tree structure is provided in Supporting Information D. Although a comprehensive review of the generated tree is beyond the scope of this work, we present here an analysis of several regions of interest in the tree. Similarly, although the opportunity exists for more detailed age estimates based on the constructed tree, this analysis is beyond the scope of the present paper.

**Figure 1.**
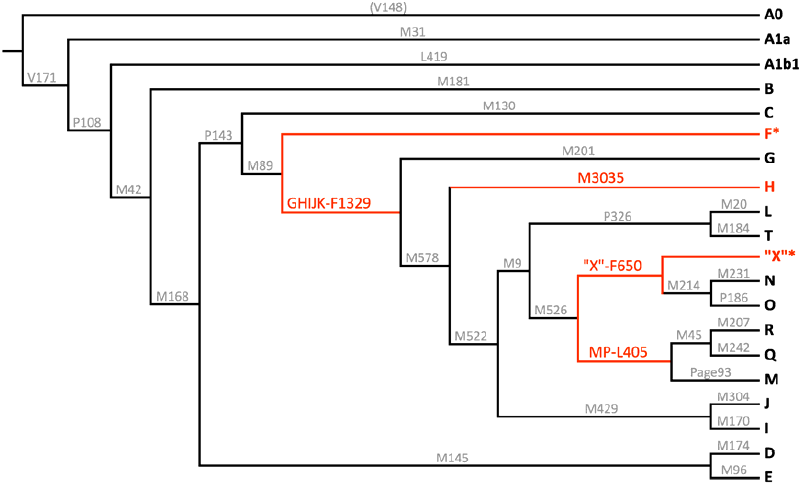
Cladogram overview of the automatically-constructed demonstration tree based on 1292-sample dataset; portions of the tree “trunk” refined through this study are highlighted; the rooting arrangement around Hg-A0 has been adjusted slightly from the automatically-generated configuration to address a rooting artifact (see main text and Supporting Information B for details)

It is noted that the dataset considered here precludes the possibility of distinguishing between mutations on the A0 branch and mutations on the A1 branch. With the chosen Hg-A0 outgroup sample, the automatically-generated phylogeny places A0 branch mutations (with inverted mutation direction) alongside A1 mutations in the main branch; and omits the A0 branch; Figure 1 has been adjusted to correct for this artifact, to be made consistent with ISOGG tree [17] configuration. We note that the consideration of genomic data for a male sample belonging to a more basal group (e.g. *Pan*/chimpanzee) would allow such ambiguities to be resolved within the presented tree generation methodology.

### Overall tree structure

Figure 2 shows the distribution of downstream branches per node of the phylogeny. The majority of nodes in the tree lead to bifurcations, however there are also many instances of 3-6 sub-branches occurring. We were particularly interested in investigating any star-like branching patterns, since these are usually associated with rapid population growth. The most extreme examples in the current phylogeny include between 8 and 18 parallel sub-branches, and are detailed in Table 1. We find evidence for star-like rapid population growth in two regions, Europe and Africa, and in two subhaplogroups, R1b-M269 and E-L576 (subgroup of E-L2).

**Figure 2.**
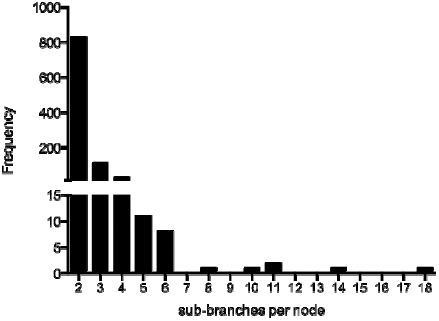
Overall morphology of the phylogeny depicted as a frequency distribution of the number of sub-branches per node

**Table 1.** Star-like phylogeny structure within the tree

| Upstream branch | Haplogroup | Parallel sub-branches | Region |
| --- | --- | --- | --- |
| DF27 | R1b-M269 | 18 | Europe |
| U181 | E-L576 | 14 | Africa |
| DF13 | R1b-M269 | 11 | Europe |
| CTS3822/M4673,<br>CTS4036/M4674,<br>CTS6126/M4676,<br>CTS7305/M4678/Z1714 | E-L576 | 11 | Africa |
| Z5943 | E-L576 | 10 | Africa |
| L2 | R1b-M269 | 8 | Europe |

Star-like phylogeny structure, indicative of rapid population expansion, was found in European and African regions of the phylogeny. In Europe much has been made of the evidence for rapid increases in population, although there is not a clear consensus on the timing(s) and locations of the expansions. In Africa the best candidate for rapid population growth is the Bantu expansion.

### Tree “trunk”

Several interesting results are apparent from the upper branches of the tree defining the major haplogroups (*i.e.* the tree “trunk”).

Firstly, the tree shows that results for the 1000 Genomes Project sample HG02040 (Kinh population) indicate a separation in SNPs defining the branch for haplogroup F (Hg-F). In particular, although HG02040 has most of the key SNPs in common with other Hg-F samples, his results indicate ancestral results for three SNPs that are found to be derived (or no-call) for all other Hg-F samples (including Hg-G, Hg-H, Hg-I, and Hg-J): 8589031 C>T (variously known as F1329, M3658, PF2622, and YSC0001299), 14367181 G>A (known as M3684 and PF2661), and 22475403 A>C (which we denote here as Z12203); there is also another SNP with similar pattern, 14237670 C>T (known as M3680 and PF2657), where the tree indicates a back-mutation in the Hg-O sample NA18994. Thus, it would appear that these three SNP mutations define a new haplogroup (which might be called haplogroup“GHIJK”) that is downstream of Hg-F and upstream of the recently-identified Hg-HIJK [3]. Due to the presence of no-calls at key sites in HG02040, further study of this sample could identify additional “approximate Hg-F” SNPs for which HG02040 is ancestral (or alternatively, there is also a remote possibility that these ancestral results in HG02040 could be shown to be spurious).

Next, the inclusion of a (relatively rare) haplogroup M sample in this tree (the Papuan, HGDP00542) appears to have some interesting implications for the phylogenetic structure around haplogroup M. In particular, the tree shows that the haplogroup M sample shares derived status with haplogroup P samples at several SNP sites, indicating the existence of an “MP” haplogroup upstream of haplogroup P and haplogroup M, and downstream of haplogroup K(xLT). (Since no Hg-S samples are considered in the present analysis, the position of Hg-S relative to this new group cannot be determined here. A similar situation exists for the smaller branches, F1, F2, and F3.) The mutations defining this “MP” branch are summarized in Table 2. With the exception of M1205, the call pattern across all the samples considered here can be explained by a single mutation (*i.e.* without apparent recurrent or back-mutations, which can be indicative of unreliability or instability). Considering the potential for no-calls in the Hg-M sample, there is the potential for additional mutations (placed at the Hg-P level in the present tree) to belong to this Hg-MP branch.

**Table 2.** Mutations defining Hg-“MP”, joining haplogroups M and P

| GRCh37 position | mutation | name(s) |
| --- | --- | --- |
| 5005053 | A>C | PF5852 |
| 14156152 | G>A | M1205 |
| 16222561 | C>T | M1221, PF5911 |
| 22222070 | A>G | PF5969 |
| 28731917 | A>C | L405, PF5990 |

Additionally, results for the sample HG03742 (Indian Telugu from the United Kingdom), when compared to other samples, indicate the existence of a haplogroup (termed Hg-“X” here until a formal nomenclature can be assigned) upstream of the present haplogroup NO and parallel to the haplogroup “MP” (discussed above) under Hg-K(xLT). The data for HG03742 indicate that he is ancestral for the haplogroup NO defining SNP mutations on the current ISOGG Y-tree: M214, P188, P192, P193, P194, and P195. However, he shares a number of mutations with the Hg-NO samples, as summarized in Table 3, with these mutations defining the Hg-“X” group. As is the case with Hg-“MP”, the position of Hg-S (along with smaller branches, F1, F2, and F3) relative to this new “X” group cannot be determined from the dataset considered here. However, it is noted that data from the Genographic Project’s Geno 2.0 microarray platform [20] considered by Morley [21] suggest that Hg-S samples will contain at least one of the Hg-“X” mutations: F650; although filtered from consideration in the tree construction approach here due to strand bias, manual analysis suggests that another mutation, F549, would be placed on Hg-“X” here and is also noted as having derived allele in Hg-S in the Morley analysis (in common with Hg-N and Hg-O samples). Thus, this is suggestive of a potential “NOS” label for the “X” branch, though it is possible that some of the other mutations on the Hg-“X” branch (besides F650 and F549) define a separate branch parallel to Hg-S. Consequently, we avoid ascribing a definitive label to this branch here.

**Table 3.** Mutations defining Hg-“X”, upstream of Hg-NO. (*italicized* entries correspond to sites with more than one mutation in the automatically-generated phylogeny)

| GRCh37 position | Mutation | name(s) |
| --- | --- | --- |
| 3909630 | *T>C* | *Z12216* |
| 7690182 | A>T | Z4842, M2308 |
| 8674808 | C>T | Z4858, M2313 |
| 9978055 | *C>A* | *Z12176* |
| 21797754 | T>C | Z4952, M2339 |
| 23208284 | G>A | CTS11667 |
| 23617006 | G>A | F650, M2346 |

The two refinements beneath haplogroup K(xLT) described here are summarized in Figure 3.

**Figure 3.**
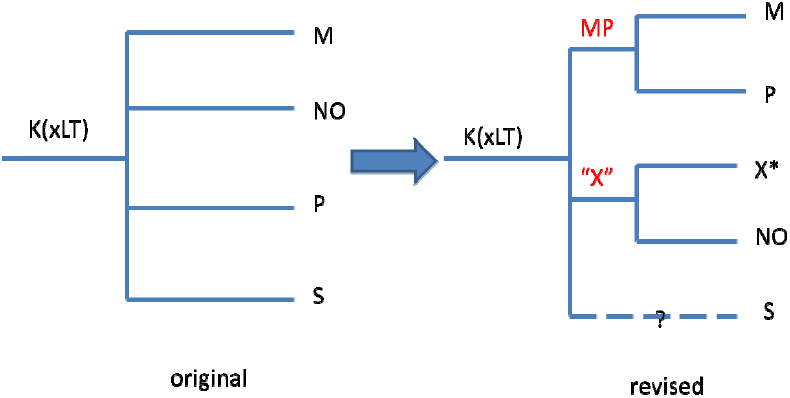
Revisions to structure beneath haplogroup K(xLT) identified with the demonstration tree

### Haplogroup A

The tree reflects the new recognition of multiple branches from the root which share the traditional designation of A, though they do not constitute a monophyletic clade. Although the very earliest branch, A00 [22], is not represented in the dataset considered here, members of A0a1 and A0b are fortunately included, and one of these was selected as outgroup sample for construction of the tree.

The Gambian A0a1 sample was found to be ancestral for all the previously known SNPs defining A0a1a and A0a1b. Because it is a singleton, a new clade cannot yet be clearly defined, but this westernmost known representative of A0a1, with its many new SNPs, is indicative of a new branch to be discovered as additional samples come to light.

The lone A0b sample is from Barbados; no African sample belonging to this extremely rare clade has yet been located, as it was first identified from a Barbadian participant in Thomas Krahn’s “Walk Through the Y” sequencing project [23] in 2012. These two A0b samples seem to share virtually the same SNPs, and are likely related.

In A1a, a clade which has in the past been lacking in internal structure, we have identified a new subclade identified by 128 new SNPs (Z11345-Z11472). Samples HG02666 and HG02645 belong to this new branch. All three A1a samples are from the Gambia.

### Haplogroup B

In haplogroup B, the seven diverse samples included in the 1000 Genomes Project, from the Gambia, Sierra Leone, Kenya and Puerto Rico allow the discovery of significant new structure at the root of the haplogroup. Three of the samples, HG02588, HG03225 and HG03376, along with the four other haplogroup B samples, share derived results at 38 SNPs, including M60, M181, and V244, which define the haplogroup in the ISOGG and other published trees. But these three samples are ancestral for two SNPs which were formerly thought to define haplogroup B, P90 and M247/P85. They are also ancestral for the SNPs that define both of the known B subgroups, B1 and B2, as shown in Table 4.

**Table 4.** Results for indicative SNPs in haplogroup B samples

| Sample | Origin | M60 (B) | M181 (B) | V244 (B) | P90 (was B) | M247 / P85 (was B) | M182 (B2) | L1391 (new B3) | M288 (B1) | M236 (B1) |
| --- | --- | --- | --- | --- | --- | --- | --- | --- | --- | --- |
| HG02588 | Gambia | T ins (Der) | C (Der) | T (Der) | C (Anc) | T (Anc) | C (Anc) | T (Der) | G (Anc) | C (Anc) |
| HG03225 | Mende, Sierra Leone | T ins (Der) | C (Der) | T (Der) | C (Anc) | T (Anc) | C (Anc) | T (Der) | G (Anc) | C (Anc) |
| HG03376 | Mende, Sierra Leone | T ins (Der) | C (Der) | No call | C (Anc) | T (Anc) | C (Anc) | T (Der) | G (Anc) | C (Anc) |
| NA19384 | Luhya, Kenya | T ins (Der) | C (Der) | T (Der) | T (Der) | C (Der) | T (Der) | G (Anc) | G (Anc) | C (Anc) |
| NA19454 | Luhya, Kenya | T ins (Der) | C (Der) | T (Der) | No call | C (Der) | T (Der) | G (Anc) | G (Anc) | C (Anc) |
| NA19043 | Luhya, Kenya | T ins (Der) | C (Der) | T (Der) | T (Der) | C (Der) | T (Der) | G (Anc) | G (Anc) | C (Anc) |
| HG00553 | Puerto Rico | T ins (Der) | C (Der) | T (Der) | T (Der) | C (Der) | T (Der) | G (Anc) | G (Anc) | C (Anc) |

These three samples’ results allow a new B3 clade to be recognized alongside B1 and B2, the members of which share novel derived SNP results at over 800 mutation sites (Z5057-Z5852) not derived in any other samples — the large number reflecting the great age of haplogroup B. Interestingly, these three 1000 Genomes Project samples also share nine more derived SNPs (L1388-L1394, L1396, and L1397), of the eleven new mutations found in the 352,776 bp tested in a 2013 “Walk Through the Y” participant from the Bahamas, another member of this clade.

Within the new B3 branch, a subclade can also be defined by the over 200 derived SNPs (Z11473- Z11671) which the two Mende samples have in common.

### Haplogroup C

The set of samples considered here has important gaps in the previously described haplogroup C geographical distribution [24, 25]. Sub-haplogroups missing from the tree include P39 (Amerindians), L1373 (central Asians), V20 (Europeans), as well as samples from countries and dependent islands southeast of the South China Sea. The chief haplogroup C subhaplogroup analyzed here is the C-M356 group of southern Asia. The tree here shows a considerable number of M356 subgroups, concentrated in the 14 P92+ Gujarati samples. Samples from East Asia carry Z1338 (or equivalents F1144, M386/CTS117), which defines a branch which, going downstream, includes Z1300 and then M407. In previous phylogenies [26], M407, one of the more recent Z1338 sub-branches, had defined this branch. Finally, the four Japanese C-P121 samples form two new branches.

### Haplogroup D

The dataset considered here lacks samples from Tibet and from among some western China groups and parts of India which together comprise a significant number of haplogroup D men [27-29]. The 1000 Genomes Project haplogroup D samples are all Japanese and within D-M55, mostly within M125, contributing about a half dozen branches. Because the samples are only from Japan, no conclusions can be made about the full geographical distribution of these new branches.

### Haplogroup E

Within Africa, the inclusion of six ethnic groups from four countries of central Africa likely provides a good cross section of haplogroup E from that area. In addition, north and central American samples, presumably as a result of the transatlantic slave trade, add to the diversity of the haplogroup E sample set. It is notable that northern and southern African populations are unrepresented in this dataset. The E in southern Africa, however, is likely there as a result of the Bantu expansion from central Africa about 3,000 years ago [30]. Within this dataset, the Luhyas of Kenya are the primary representatives of the eastward Bantu expansion [31] though their migration was relatively short geographically. The Mende of Sierra Leone are also migrants, in this case from the east or north sometime over the last 2,000 years [31]. Gambians, Esans, Yorubas and Maasai do not have a history of long migrations to their current residences.

A large number of haplogroup E samples are available in this dataset and can be divided into six sections.

*<u>E-M33:</u>* The men in this dataset within M33 are seen to be overwhelmingly West African.

*<u>E-V95:</u>* The Mende group members cluster heavily within the general V95 subgroup; the Gambians within L485. Otherwise there is considerable mixing within V95 of West Africans and descendants of slaves in the Americas. A large number of new subgroups are recognizable within V95 on the tree, with special mention provided here for the large M191 and U175 subgroups of V95.

*<u>E-M191:</u>* Over 60 new M191 subgroups are identified in the tree here. The Maasai and Gambian populations are un-/under-represented among the M191 samples. Among M191 samples, almost all fall into either subgroups of west African or Luhya ancestry. A single Mbuti Pygmy sample does not cluster with the others. One M191 subgroup is predominantly Esan from Nigeria, and joined as nearest relatives by Afro-Caribbean populations.

*<u>E-U175:</u>* Almost 50 new U175 subgroups are identified in the tree. The sub-branches in this part of the phylogeny overwhelmingly cluster by source population. The U175 men from the Americas are intermixed throughout.

*<u>E-M215:</u>* Almost a dozen M215 subgroups not listed in the ISOGG Y-tree [17] are shown in the generated phylogeny. All the available M215 samples are within the M35.1 branch. They fall almost equally into either the Z827 or V68/L18 subbranches of M35.1. The Z827 men are predominantly from Hispanic areas, *i.e.* Iberia or the Americas. There is a separate Luhya, Kenya, branch off of Z830 within Z827, and another Z827 branch within L19 has three samples of which a Gambian and Afro-Caribbean sample form the majority. The V68/L18 branch also has a distinctive Luhya branch, and, overall, samples from Tuscany in Italy are more common than in the Z827 branch. Hispanics are also well represented in V68/L18.

*<u>E-M75:</u>* Three new E-M75 subgroups are identified in the phylogeny. In addition to a distinct M200+ subgroup in the Maasai of Kenya, there are a few West Africans and a single sample from among the Dinka of Sudan.

### Haplogroup G

Prior studies have shown haplogroup G is most common in the Caucasus mountain regions [32], which were not sampled by the 1000 Genomes Project. The haplogroup G samples in the dataset considered here encompass almost all the major G branches otherwise known earlier through detailed testing, including subgroups common in the Caucasus. There are two important omissions included in the ISOGG Y-tree [17]; (a) the G-L177 subgroup which has many Sardinian G men who share SNPs equivalent to L177, and (b) two subgroups of G1, L1324 and L830.

The Y tree here shows the phylogenetic relationships of G-Z724 and G-Z1903, comprising an important branch most common in Europe, particularly Sardinia [2], but not discussed in prior studies. In addition, newly identified is a branch (CTS4367, L1259, PF2970) joining the PF3146 and L30 subgroups. Also within haplogroup G, the Punjabi G-L166 sample, HG02681, here shares four L166-specific SNPs with the 5,300-year-old G-L166 Iceman mummy Ötzi, found in the Italian part of the Ötztal Alps [33] (data from ERP001144), and the same number with a subset of G-L91 samples from Corsica, Sardinia or Tuscany, inferred to be also G-L166 [2]. These SNPs (PF3178, Z6134, Z6213 and Z6287) are ancestral in the Puerto Rican L91+, L166- sample, HG01311.

### Haplogroup H

The tree here provides considerable revision of the previous 2008 haplogroup H tree. The revised H tree is defined by M3035, which arose earlier than its two principal subgroups, the previously identified M69 subgroup and a new subgroup (samples are labeled in supporting information per existing ISOGG criteria as haplogroup F-M89). A recent study reported that F3 is also a branch of M3035 [34]. The present study lacks F3-M282 samples.

Haplogroup H is heavily concentrated in southern Asia, with a secondary concentration among the Roma population of Europe and in isolates in surrounding regions [35, 36]. The new H branch is a mix of south Asian populations sampled: Telegus, Sri Lankans and a single Punjabi. Ten new subgroups are identified within this new branch. There is intermixing of Telegus and Sri Lankans within subgroups. Among the traditional M69 H men, they are overwhelmingly M52 men and come from a wider variety of H men than in the new branch. Within M69, Gujuratis of western India and Bengalis of Bangladesh and an occasional Pakistani Punjab sample are found. About 30 M69 subgroups are recognized in the tree here. None of the major clusters is confined to a single ethnic group. A new upstream subgroup is identified for the mutation Apt. The samples are a mix of Bengalis, Sri Lankans and Telegus, and new subgrouping parallel to Apt is identified.

### Haplogroup J2a-M410

Haplogroup J2a is classically described as the main vector of the Neolithic expansion of agriculture to Europe and western Asia [37, 38] and has more recently also been connected to metallurgy networks in Eneolithic/Copper Age [39, 40].The 1292-sample dataset employed here is able to confirm most of the haplogroup J2a phylogeny as recognized by ISOGG [17] and previous publications, and also shows the most important new sub-nodes of J2a-M410 partially published in 2013, based on Sardinian sample results [2], analysis of new Genographic Project Geno 2.0 results [20, 21] and a minimal reference phylogeny [34]. This in spite of the absence of data from the Middle East, in which the northern part is a center of J2 frequency and diversity [35, 41-44]. SNP markers with names in the range Z6046 through Z8120 were identified and provisionally placed with the data for samples considered here. The J2a structure from the computer-generated demonstration tree is shown in Figure 4.

**Figure 4.**
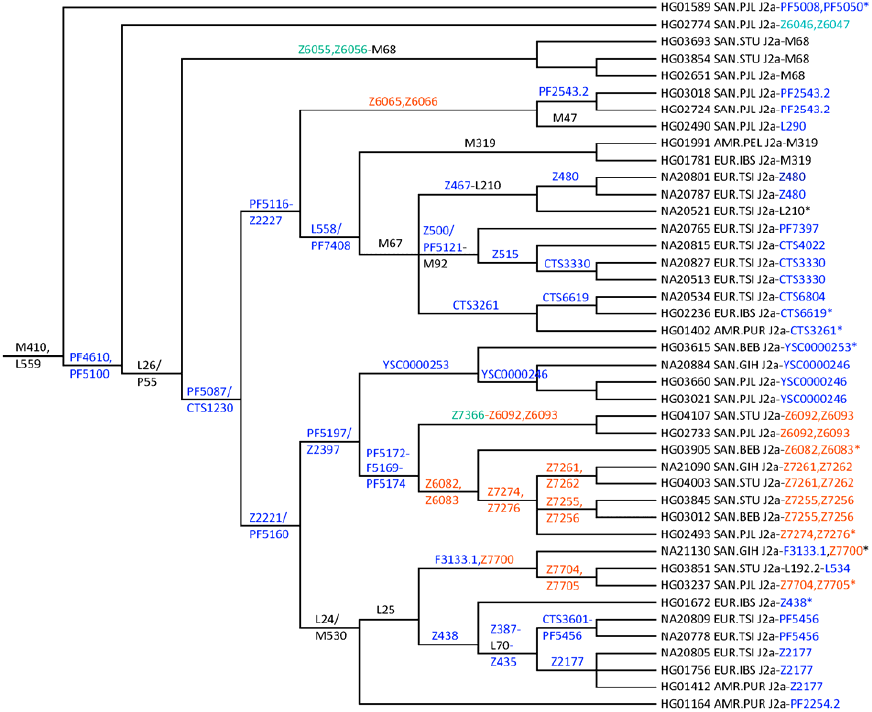
Structure for haplogroup J2a in the demonstration tree. Clades with only colored markers are not present in the latest ISOGG Y-tree. Blue color: previously identified (partly placed) markers; green color: markers previously identified in Sardinians; red color: markers identified and provisionally placed in this study.

Data for additional samples from the aforementioned Sardinian [2] and Geno 2.0 [21] studies are considered in the following discussion to supplement the results that are apparent from the tree generated here.

J2a-PF4610* is revealed by a Punjabi sample and independently confirmed by seven Sardinian SNP results. By comparisons, this ancient lineage is expected to be defined by Z6046 and its equivalents.

J2a-PF5008* has one Punjabi sample — unfortunately with a no-read for L581 — which thus could be positive or downstream one clade by comparison with ten Sardinian samples. Further substantial new substructure is only revealed by those Sardinian samples and Geno 2.0 results with the defining SNPs PF7381, PF7384, PF4993 and equivalents. Together with J2a-Z6046, the proposed J2a-PF5008 haplogroup should be able to reveal the affiliation of most J2a* samples in previous research.

J2a-M68 has an Eastern distribution, with two Tamil and one Punjabi samples, and is believed to lie within a haplogroup defined by SNP Z6055 and equivalents, by comparison with two Sardinian samples. This proposed upstream haplogroup J2a-Z6055 could be interesting in future studies of the remaining J2a-L26* haplotype clusters.

J2a-PF5087/CTS1230 defines a new macro-haplogroup downstream of J2a-L26, which includes prominent J2a-lineages like M67, M319, L24/M530, etc.

J2a-Z2221/PF5160 unites the known and widespread haplogroup L24/M530 [43] with a new clade, defined by PF5197/Z2397, below PF5087/CTS1230.

J2a-L24/M530 is subdivided into the known L25 and the new minor clade PF2254 as revealed by a Puerto Rican sample and confirmed by one Geno 2.0 result. L25 has two subclades Z438, with five west Mediterranean samples and one from Puerto Rico, and F3133 with three Asian samples within J2a. An important subclade of Z438 is Z387, until now recognized by the STR marker value DYS445≤7 value [17]. With only one Iberian sample, PR329 defines a (probably minor) subclade of Z438. Z387 has the known subclade L70 which has its own subclade, Z435, further subdivided into CTS3601 and Z2177. From Geno 2.0 data comparison, CTS3601 should have substantial substructure defined by PF5456, here positive in two Tuscan samples. The J2a-F3133 haplogroup could be probably more reliably described by Z7700 and equivalents. Its subclade, Z7704, has the known subclades L192 and L534 below it. Comparison with Sardinian samples and Geno 2.0 results indicates that PF5294, a major subclade of L25, along with its subclades (PF4888, L243, PF5366, and others) are missing from the 1292-sample dataset considered here.

J2a-PF5197/Z2397 is a new haplogroup with 13 samples belonging to two subclades. The most important is J2a- PF5174, with three sequences from Tamil, two Bengali and two Punjabi as well as one Gujarati. When 13 Sardinian samples are also considered, a distribution from South Asia to Europe is apparent. Interesting substructure could be revealed by the proposed sub-haplogroups defined by Z6082 and Z7366. Z6082 has the subclade Z7274 further subdivided into Z7255 and Z7261. Z7366, by comparison with three Sardinians, is subdivided in Z6092 and PR2128. Ten Sardinians and Geno 2.0 results reveal substructure of PF5174 defined by PF5177. Geno 2.0 results also indicate structure upstream of PF5174 with PF5172 and PF5169 and equivalents. PF5169 should include the known ISOGG haplogroup J2a-L198 and together with other Geno 2.0 results defines another J2a. The second subclade of J2a-PF5197/Z2397 is defined by YSC0000253 (among others) with two Gujarati and one Punjabi samples. A major subclade is defined by YSC0000246.

J2a-PF5116, Z2227 sub of PF5087/CTS1230 unites known and new haplogroups with expected expansion mainly to Europe. Downstream is L558/PF5121 uniting ten M67 and two M319 samples from Iberia and Peru. Also downstream is Z6065 etc. uniting one M47 sample from Punjab and an unknown lineage defined by another two Punjabi samples with expected low diversity. Except for M67 there are no Sardinian samples for comparison. Geno 2.0 data however reveals that PF5116 could be upstream of Z2227.

J2a-M67 is shown by three subgroups. Three Tuscan samples are positive for the L210 SNP. One clear subgroup is defined by Z480 etc. Taking into account Geno 2.0 results, a group upstream to L210 defined by Z467 is revealed. Four Tuscans reveal the substructure under M92: one group should be defined by Z515 and the subgroups CTS3330 confirmed by three Sardinians and CTS4022. The remaining sample should be CTS2906, confirmed by three Sardinians. From comparison with another three Sardinians, CTS12 should describe an upstream group below M92. Another four Sardinians show that upstream of M92 there is a haplogroup Z500 with another subgroup PF7394. Two further samples from the Western Mediterranean and one from Puerto Rico show the third sub-lineage of M67 as being CTS3261. One subclade, defined by CTS6619, is found to be further subdivided by CTS6804 by comparing with Sardinian and Geno 2.0 results. Comparison with Sardinian sample set suggests that not all M67* samples could be addressed with the groups described here.

### Haplogroup J2b-M12

Haplogroup J2b-M12 had little representation in this study; however when correlated with data from other publicly available genomes and haplogroup projects, some observations can be made. M12 now sub-divides into three branches defined by M205, M241, and Z2453.

J2b1-M205 exists in low frequencies throughout Europe, the Mediterranean, and the Middle East. J2b2-M241 is distributed throughout Europe and the Middle East, as well as west and south Asia. It divides further into two sub-branches L283 and Z2432.

J2b2a-L283 was discovered by Family Tree DNA through its “Walk Through The Y” program, and is predominantly Middle-Eastern, Mediterranean and European. The M12/M241 frequency peak in the Balkan Peninsula and Italy observed by Semino *et al.* [37] and Cruciani *et al.* [45], may instead belong to sub-clade L283. A recent Z631 sub-branch expansion from east to west through the heart of Europe to the UK along with presence in Italy and Spain might be associated with Roman expansion using mercenaries and slaves acquired in the Balkans.

J2b2b-Z2432 is a significant new clade which exists primarily in south-west Asia, particularly India and surrounding regions.

Although Z2453 is seen as a singleton in this tree, present in the 1000 Genomes Project sample, NA20588, from Tuscany, Italy, comparison with Personal Genome Project data [46] identifies J2b3-Z2453 as a new clade.

Additional polymorphisms previously listed below M241 including M99, M280, M321, and P84 [26, 47-49] were not observed in this study. A notable cluster below J-M241 defined by DYS455=8 [50] also does not yet have sufficient genotyping available to completely determine its relationship with the above polymorphisms, and as of the time of this writing, has been removed from the ISOGG Y-tree [17].

### Haplogroup L

The world population of haplogroup L is concentrated in southern Asia with small percentages elsewhere [35]. This origin is reflected in the 1000 Genomes Project haplogroup L samples which all come from that area. The samples, however, all fall either in the M27 or M357 areas of the tree. The tree identifies almost 14 new subgroups under M27 and five in the M357 area. None of the subgroups is composed of just a single population group.

### Haplogroup N

Almost all haplogroup N men are found primarily in a geographical belt extending from eastern Siberia and the Siberian southern border area to Finland and the Baltic nations [51, 52]. A small number of samples from Beijing, China, and a few from southern Asia comprise two new branches under general N-M231. The rest of the N samples in 1000 Genomes Project come from VL29 or Z1936 men from the Helsinki, Finland, sampling site. Central Asia and Siberia were not included in the project. The tree shows two new subgroups under VL29. Within the combined Z1936/Z1925 branch, Z1940 accounts for most of the samples, with five apparent new subgroups. There is a parallel branch to Z1940 containing a few samples, and four new subgroups from a branch just downstream from Z1936/Z1925.

### Haplogroup Q

Haplogroup Q is found overwhelmingly either in eastern Russia, northern China or among Amerindians of the Americas and in small percentages elsewhere [53-55]. In the 1000 Genomes Project, the haplogroup Q samples are only from Mexican, Colombian and Peruvian men of presumed Amerindian ancestry, together with a few samples from southern Asia. Also included is a sample from an Amazonian Karitiana Amerindian. The Amerindians are all within Q-M3. The Amerindian samples provide almost 20 new subgroups for the tree, and the Asian men provide a few subgroups under either L472 or L275. Clustering by country is noted among the Mexicans, Peruvians (two subgroups) and the small number of Colombians. The Karitania sample has 154 singleton Y mutations. The Mexican-Colombian-Peruvian samples typically have fewer, but a few samples have similar numbers of singletons as the Amazonian. Higher-coverage sequencing of the Amazonian sample partially explains its results. The most recent common male ancestor of some of the men with higher numbers of singletons may thus have lived as far back as the estimated time of the first settlement of the Americas (13 to 20 kya [56] depending on the study) or possibly earlier. In contrast, the Peruvian Q-M120 sample HG01944 has about half as many singletons as the Amazonian one, and its M120 status is unexpected if the sample might be from an Amerindian. Q-M120 is usually found in eastern Asia, and in this tree, it is otherwise only found in Vietnamese samples. Q-M3 is the mutation traditionally associated with Q Amerindians and with most Peruvian samples here.

### Haplogroup T

Haplogroup T has mostly a Mediterranean Sea distribution with some prevalence in a few Jewish groups [2, 38, 57]. There are only eight T samples in the 1000 Genomes Project, all from the western Mediterranean or descendants of migrants from there to the Americas. The tree here identifies new subgroups downstream from P77 and under L446.

## Conclusions

The approach described here has been demonstrated to construct a reliable, high-resolution Y-chromosome phylogeny based on (primarily low-coverage) short-read sequencing datasets. The *a priori* nature of the approach makes it readily generalizable to other applications and flexible with regard to errors and omissions in the current state of knowledge of the Y-tree; at the same time, this aspect means that the approach becomes very computationally expensive (in terms of both CPU time and memory) as the number of samples considered grows, so, when considering larger datasets, careful sample selection will likely prove useful for efficiently studying particular regions or features of the Y-tree.

It appears that iterative reweighting plays an important role in the robustness of this tree generation approach, by effectively removing many spurious variants and unstable sites and thus taking some of the burden off of variant filtering, allowing for generally less restrictive non-position-based filters to be used. These non-position-based filters, in conjunction with the iterative reweighting process, can effectively adapt to the underlying quality of the data and their propensity to suggest false positive variants. Although this approach will require more careful consideration (than with position-based filtering) when applied to studies that depend on region size (e.g. mutation rate and haplogroup age estimation investigations), it should be more flexible with regard to underlying data quality and also appears to be generally less restrictive in terms of excluding phylogenetically informative variants (as noted above), allowing a higher-resolution phylogenies to be obtained.

Iterative reweighting, being the most unique aspect to the approach described here, appears worthy of further study and refinement. For example, the success of this approach described here, aided significantly through the use of iterative reweighting, suggests the possibility for improving robustness and speed through integration of a similar sort of dynamic reweighting directly into the underlying phylogenetic tree generation algorithms in software packages such as PHYLIP.

Furthermore, the tree-generation procedure described here, particularly the novel iterative reweighting approach, may find use in other areas of molecular phylogenetics, including the generation of *a priori* phylogenies for viral, bacterial, and mitochondrial systems.

## Acknowledgements

The authors gratefully acknowledge helpful discussions with Steve Fix and Richard Rocca during refinement of the tree generation methodology and application to the 1292-sample dataset considered here. The authors also acknowledge helpful discussions with Joe Felsenstein related to memory management in PHYLIP. The authors would like to thank David Reynolds for his work on compiling and maintaining the database which has recently become the ISOGG SNP Compendium. We gratefully acknowledge Thomas Krahn for sharing details of “Walk Through the Y” findings with several of the authors. Finally, the authors acknowledge helpful discussions with Justin Loe regarding computational phylogeny considerations.

## Supporting information

Supporting Information A: Implementation details

### Variant calling

Variants are called from BAM files using the following “mpileup” command from *samtools* v. 0.1.19:

$ SAM_HOME/samtools mpileup -C50 -F 0.5 -ugpf $REF_FILE [bam file list] |
$ SAM_HOME/bcftools/bcftools view -vcgm0.99 - > [output VCF file]

The reference sequence from the 1000 Genomes Project is used: http://ftp.1000genomes.ebi.ac.uk/vol1/ftp/technical/reference/human_g1k_v37.fasta.gz

### Genotype refinement

Genotype (GT) calls are refined using a python script that reads the VCF file and produces a VCF file with refined genotype calls. The algorithm for calling based on each set of genotype likelihoods (PL values) is summarized as follows:

*-look for zero in PL values corresponding to homozygous calls*

*-if 0 is found at one of these sites, check that all the other PL values corresponding to homozygous calls are at least 10 (phred scaled likelihood) and all other PL values corresponding to heterozygous calls are at least 3; if these criteria are met, return the genotype call corresponding to the 0 PL value; otherwise, continue*
*-look for zero in PL values corresponding to heterozygous calls*

*-if 0 is found at one of these sites, check that all the other PL values corresponding to homozygous calls are at least 2 (phred scaled likelihood); if these criteria are met, return a heterozygous call, “./.”; otherwise, continue*
*-if none of the above criteria result in call, return a no-call (“.”)*

### Variant filtering

Annotation with the *vcf-annotate* utility of *vcftools 0.1.10* is performed (prior to the aforementioned genotype refinement) with the following command-line options:

*-f HetNew.filt -f +/D=7800/q=13*

The q=13 option requires a minimum root-mean-squared (phred-scaled) mapping quality of 13 (“MinMQ” filter); D=7800 imposes an upper limit on depth of coverage of 7800 reads across all 1292 samples (“MaxDP” filter). We have found that a cutoff of 6 reads is a reasonable heuristic for the low-coverage 1000 Genomes Project samples, and this cutoff depth is set fractionally higher here to account for the small number of samples with greater average coverage depth. The “+” option requests all the other default filters, including considerations like strand bias.

The specification of custom filter file (“HetNew.filt”) in the command-line options above also applies the following two filters:

“Het”: G3 corresponding to heterozygous 0/1 genotype exceeds 0.025

“Het2”: G3 corresponding to heterozygous 0/1 genotype exceeds half of G3 corresponding to 1/1 genotype

(G3 corresponds to the maximum likelihood estimate of genotype frequencies reported for many of the variant sites by samtools.)

Various other custom filters are applied using a custom python script, following the aforementioned genotype refinement; the filters applied in this step are summarized below.

“LongIndel”: sites with REF or ALT alleles with more than a threshold length are marked with this filter (with the hypothesis that “longer” INDEL allele representations will be more mutationally-unstable, and potentially also less reliable. A threshold of 4 bp is used here.

“Het3”: sites with ratio of ‘./.’ calls to (total - ‘.’) greater than a threshold are marked with this filter. A threshold of 0.025 is used here.

“FewALT”: sites with fewer than threshold value number ALT calls (according to the refined GT call) are marked with this filter. A threshold of 2 is used to exclude “singletons” whereas a threshold of 1 is used to consider these “singletons”.

“HighNC”: sites with high ratio of ‘.’ + ‘./.’ calls to total samples are marked with this filter (high no-call rate). A threshold of 0.381966, based on the golden ratio, is used here.

“MNP”: sites with MNPs (multiple nucleotide polymorphisms) of size larger than some threshold are marked with this filter. A threshold of 1 bp is used here, effectively excluding all MNPs

“ManyALT”: sites with a high number of ALT alleles, greater than some threshold, are marked with this filter. A threshold of 2 is used here.

“Nallele”: sites with REF or ALT containing N are marked with this filter

“Het4”: sites with ratio of Het (“./.”) to ALT calls greater than the threshold value are marked with this filter. We have found that using this filter can be overly restrictive, excluding some genuine SNPs, and have found iterative reweighting, on the other hand, to serve as a more effective alternative, allowing for generation of reliable trees while retaining these sites. Consequently, the threshold is set sufficiently high here to effectively disable this filter.

### Maximum parsimony tree generation

The *pars* program (discrete character parsimony algorithm) within PHYLIP 3.69 (with modification to memory management for ancestral state reconstruction, as described in text) is used for maximum parsimony tree generation. Critically, options to print out steps at each site and to print the character at all nodes of the tree are turned on.

For the example tree shown here, the “less thorough” search option is specified, with one tree saved. A random seed of 9 is used, with 1 jumble. The sample HG01890, known from previous investigations to belong to Hg-A0, is specified as outgroup.

### Iterative reweighting

The number of state changes (mutations) of each site throughout the tree is tallied and used to determine weighting for the next tree generation iteration. The weighting schedule summarized below is used:

| Reweighting iteration | 1 | 2 | 3 | 4 |
| --- | --- | --- | --- | --- |
| Weights determined based on mutation count in: | Tree 0 | Tree 1 | Tree 2 | Tree 3 |
| Weights determined in this iteration are used to generate: | Tree 1 | Tree 2 | Tree 3 | Tree 4 |
| Weighting function, $W(M)$ ($M > 1$) | $\frac{35}{M-1}$ | $\frac{35}{(M-1)^2}$ | $35e^{-2(M-2)}$ | $35e^{-2(M-2)}$ |
| Zero weight for nearby mutations on same branch? | no | no | no | yes |

“Tree 0” is generated without weighting. The weighting function in the table above is applied to determine an integer weight (determined with rounding) from 0 to 35 (inclusive) based on the mutation count, M. For M=0 and M=1, the full weight of 35 is used, So, for example, with the first reweighting iteration, the following weights are used:

| M (mutation count) | W(M) (site weight) |
| --- | --- |
| M=0, M=1, or M=2 | 35 |
| M=3 | 18 |
| M=4 | 12 |
| M=5 | 9 |
| M=6 | 7 |
| M=7 | 6 |
| M=8 | 5 |
| 9≤M≤11 | 4 |
| 12≤M≤15 | 3 |
| 16≤M≤24 | 2 |
| 25≤M≤71 | 1 |
| M≥72 | 0 |

The final (4th) reweighting iteration uses the same weighting function based on mutation count as the 3rd reweighting iteration, but also imposes an additional filter that reweights sites to zero when they are within 100 bp of another site with a mutation on the same branch.

### Post-processing of tree-construction results

For the example tree discussed here, the mutations are annotated with names based on the version of the ISOGG SNP Compendium (part of the ISOGG Y-Tree resource) last updated September 15, 2013; this database includes 47,680 variants with mutation information. Additionally, a custom SNP name supplement, consisting mostly of names assigned through this work, was used to supplement the ISOGG SNP Compendium annotations. The full list of variants named with Z-prefix is maintained in a collaborative Google Document accessible via https://sites.google.com/site/yanalysis/.

Supporting Information B: Tree generation result files for the 1292-sample example

An ASCII tree file, a NEWICK tree file, and an “annotations” report localizing mutations to individual branches are included as supporting files. It is noted that the ASCII tree file is not “to-scale”. Population and haplogroup classification (based on the ISOGG tree) have been added to the ASCII tree as annotations in a post-processing step. Branch lengths in the NEWICK file do not include “singletons” for the terminal branches. In the annotations file, an asterisk (“*”) indicates that the placement of the mutation in the tree is uncertain due to no-calls and could be higher in the tree. For the rare cases of sites with multiple non-reference alleles, there is the possibility for inaccuracies in the ancestral allele in the provided mutation information. As discussed in the main text, it is noted that the use of the Hg-A0 sample, HG01890, generates here a rooting arrangement resulting in mutations associated with Hg-A0 being shown (with reversed polarity) alongside Hg-A1 mutations on the branch from node 270 to node 416. Aside from this single outgroup/rooting artifact, the generated tree agrees with the traditional arrangement. Finally, the tree generation approach will sporadically identify “empty” branches in which no mutations can be solidly localized to this particular branch, though there is the potential for a distinct branch to exist here due to no-calls in particular samples in the underlying data; in analyzing these cases with effectively zero branch length, it seems best to assume (as a null hypothesis to maximize parsimony) that such “empty” branches do not exist as a distinct part of the phylogenetic structure.

Supporting Information C: List of BAM filenames used in 1292-sample tree

A list of BAM filenames used in this study is included as a supporting file. The files from 1000 Genomes Project contain additional details including sequencing platform and alignment date.

Supporting Information D: Cladogram for the 1292-sample tree

An overview of the generated tree structure, including sample names and labels for major haplogroups, is provided in PDF format.

